# Comparison of volumetric dynamic optical coherence tomography with biological methods for evaluation of radiation effects in prostate tumor spheroids

**DOI:** 10.1101/2025.11.08.687368

**Authors:** Steph Swanson, Keyu Chen, Elahe Cheraghi, Ernest Osei, Kostadinka Bizheva

**Affiliations:** Department of Physics and Astronomy, University of Waterloo, Waterloo, Ontario, N2L 3G1, Canada; Department of Medical Physics, Waterloo Regional Health Network, Kitchener, ON, N2G 1G3, Canada; Department of Clinical Studies, University of Guelph, Guelph, Ontario, N1G 2W1, Canada; School of Optometry and Vision Sciences, University of Waterloo, Waterloo, Ontario, N2L 3G1, Canada; Systems Design Engineering Department, University of Waterloo, Waterloo, Ontario, N2L 3G1, Canada

## Abstract

**Significance:** 3D tumor spheroids are more physiologically representative of *in vivo* patient tumors compared to 2D monolayer culture. However, their 3D nature challenges the use of conventional biological techniques like proliferation assays, fluorescence microscopy, and the clonogenic assay, which is the gold standard method for assessing cell survival following radiation. However, clonogenic assay requires spheroid disaggregation.

**Aim:** Non-invasive volumetric imaging with dynamic optical coherence tomography (dOCT) enables cellular activity to be visualized with spatial resolution within 3D tumor spheroids. Cellular activity observed via dOCT in irradiated prostate tumor spheroids was quantified for comparison with conventional biological techniques.

**Approach:** A Varian TrueBeam linear accelerator was used to irradiate spheroid and monolayer cultures with a 6 MV beam. Cellular activity was estimated from dOCT images generated via frequency banding and compared to clonogenic assay, proliferation assay, fluorescence microscopy, and 3D cell simulation.

**Results:** Prostate cancer cells cultured as spheroids demonstrated improved radio-resistance via clonogenic assay compared to monolayer culture. The dOCT method demonstrated quantitative and qualitative agreement with proliferation assay and fluorescence microscopy, respectively.

**Conclusions:** A longer duration of repeated dOCT measurement in tumor spheroids following radiation treatment could offer a non-invasive alternative to the clonogenic assay.

## 1 Introduction

Prostate cancer is the most prevalent cancer in North American males [1, 2] and is commonly treated with radiation therapy [3]. Clinically, modern radiation treatments are delivered through fractionation schemes that leverage distinct radiobiological behavior exhibited by different tumor types and their surrounding tissues [4–6]. Our foundational radiobiological understanding of cellular responses to radiation was formed through conventional 2D monolayer cell culture methods [7, 8]. In 1956, Puck and Marcus [9] irradiated mammalian cells cultured in 2D monolayers and published the first radiation survival curve that fit cell survival to radiation dose using a colony-forming or clonogenic assay. Since then, the clonogenic assay has remained the gold standard for measuring radiation survival [10], in part because it informs the extensively validated linear-quadratic (LQ) model [11, 12]. In the LQ model, the linear *α* and quadratic *β* parameters mechanistically describe cell killing due to one or two radiation tracks, respectively [13]. The *α/β* ratios obtained from conventional 2D monolayer cell culture experiments continue to guide clinical decision-making [7].

However, a growing body of evidence in recent decades has demonstrated that *in vitro* cells cultured in 2D monolayers on flat, hard surfaces do not accurately replicate the behavior of *in vivo* tumors [14, 15]. In contrast, cancer cells cultured in small 3D aggregates called tumor spheroids exhibit more physiologically relevant behavior due to increased cell-to-cell contact and diffusion-limited distributions of oxygen and nutrients [16–18]. This heterogeneous distribution leads to the formation of a hypoxic spheroid core of non-proliferative or quiescent cells [15]. It has been widely observed that radiation survival differs significantly between *in vitro* cancer cells cultured in 2D monolayers and those cultured in 3D spheroids [19–21].

Cell culture geometry is suspected to influence the effects of radiation through biological and physical processes related to nutrient and waste transport, resource competition, and mechanical forces exchanged between cancer cells and their microenvironment [19, 22, 23]. Mathematical *in silico* models that simulate these biological and physical processes have become increasingly prevalent as tools to elucidate the mechanisms underlying tumor growth and treatment response [24, 25]. In particular, *in silico* methods have been utilized to investigate the mechanical and molecular characteristics of tumor spheroids [26]. Conventional *in silico* approaches, such as continuum models described by partial differential equations (PDEs), are mathematically well-characterized and computationally efficient; however, these models struggle to accurately capture the inherent cellular heterogeneity and diversity present within patient tumors and tumor spheroids [27]. In contrast, discrete *in silico* methods such as agent-based models address these limitations by simulating each cancer cell as an individual “agent” that evolves through time by responding to its local environment, including neighboring cells and nutrient availability [24, 28]. The utility of *in silico* models for falsifying hypotheses and generating novel insights depends critically upon the accuracy of their parameter estimates, which are derived from *in vitro* and/or *in vivo* measurements [29].

However, the 3D structure of tumor spheroids poses challenges for measurement by conventional biological techniques. Clonogenic assay can be adapted for evaluating radiation survival in spheroid cultures by disaggregating the 3D cellular structures into a cell suspension for 2D plating and longitudinal observation of colony formation. Nevertheless, the geometry of cell culture following radiation treatment itself influences cellular survival and response [20]. Alternative methods, such as proliferation assays, avoid the need for spheroid disaggregation and have also been utilized to estimate clonogenic radiation survival [30]. For instance, the Alamar Blue (AB) proliferation assay is a common technique that measures cellular metabolism by detecting the reduction of non-fluorescent, blue resazurin to fluorescent pink resorufin [31]. Although the AB proliferation assay has been adapted for spheroid cultures [32], the fluorescence measurement reflects only an averaged response across the entire cell population within the spheroid and does not distinguish the treatment effect in the hypoxic core from the spheroid periphery. Fluorescence microscopy (FM) is widely employed to evaluate spheroids with spatial resolution [20, 33, 34]; however, these assessments remain qualitative rather than quantitative.

In contrast, optical coherence tomography (OCT) enables the quantitative evaluation of 3D tumor spheroids through non-invasive, high-resolution volumetric imaging [35, 36]. Recently, dynamic OCT (dOCT) methods have been developed, which involve repeatedly acquiring OCT images at each spatial location to estimate cellular activity from temporal OCT intensity signal fluctuations [37]. Previously, dOCT has been employed to investigate spheroids [38–41] as well as other types of 3D cell cultures [41–44]. Quantitative analysis of tumor spheroids using dOCT has been conducted following chemotherapy treatment [38, 39] and exposure to perfluorooctanoic acid [41]; however, the effects of radiation treatment have not yet been explored with this method. Although the clonogenic assay remains the gold standard for measuring cell survival after radiation exposure, there has been sustained interest in adapting alternative methods, such as proliferation assays, to circumvent the long experiment duration and labor-intensive colony counting required by clonogenic assays [30, 45, 46]. Given prior suggestions that dOCT can quantify cellular activity and viability [38, 40], it may also provide another feasible alternative to the clonogenic assay. In this study, we employed a previously described high-speed spectral-domain line-field (LF) dOCT system [47] to image prostate tumor spheroids after radiation treatment and performed qualitative and quantitative comparisons with established biological methods and computational simulation.

## 2 Methods

### 2.1 Prostate tumor spheroids

PC3 cells (human prostate adenocarcinoma) were cultured in RPMI 1640 (Sigma-Aldrich, USA) supplemented with 1% penicillin (Thermo Fisher Scientific, USA) and 10% fetal bovine serum (Thermo Fisher Scientific, USA). Cells were incubated at 37°C with 5% CO_2_ in a humidified atmosphere. Monolayer cell culture was plated in T-25 culture flasks. Prostate tumor spheroids were seeded and cultured in a 3D Petri Dish (MicroTissues Inc., USA) composed of 2% agarose gel. Each agarose gel contained a 9 × 9 array of wells and was seeded to form spheroids composed of approximately 1,000 cells in each well (Fig. 1A).

**Figure 1.**
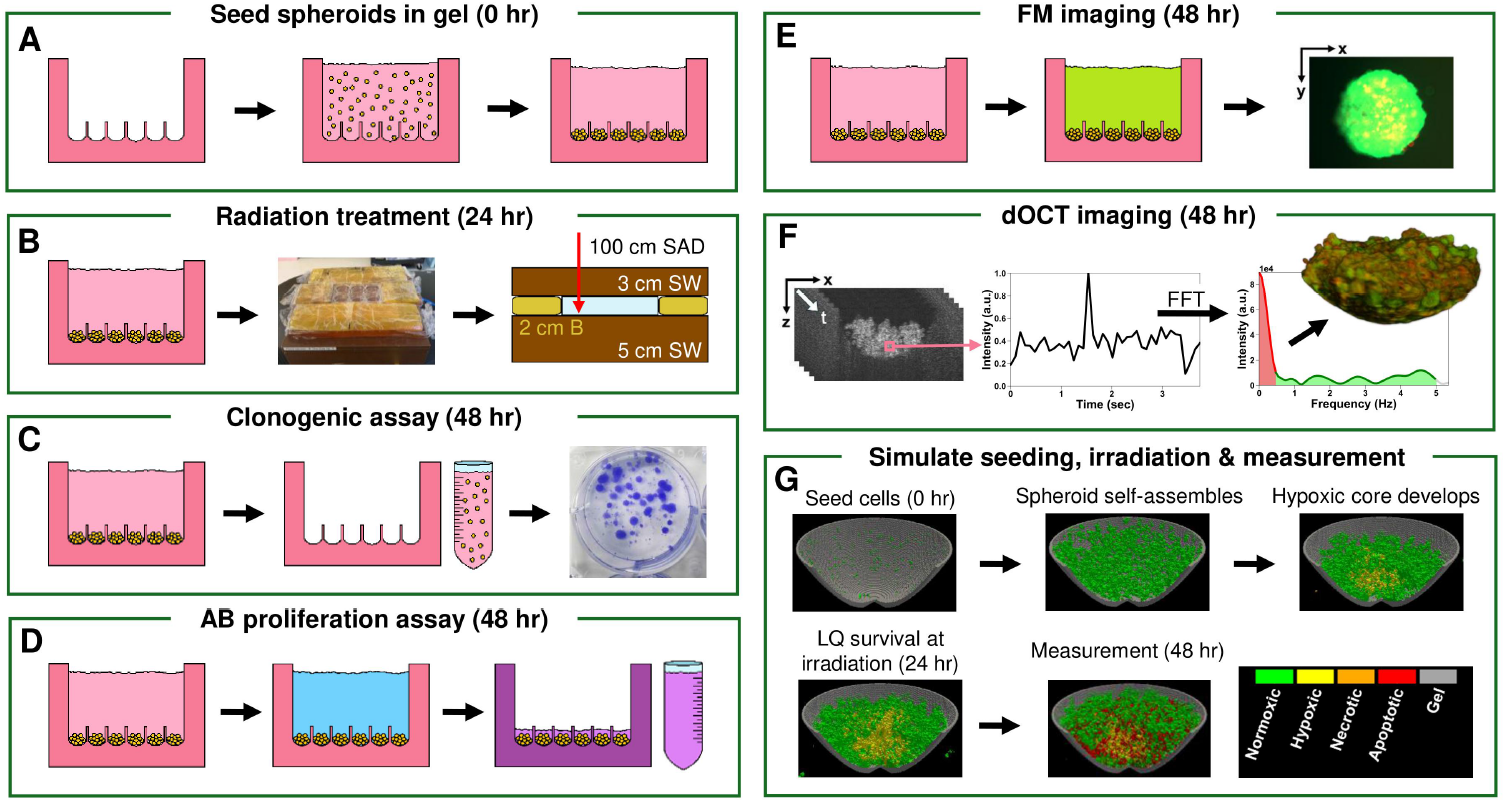
Schematic of experimental and simulated spheroid seeding, irradiation, and measurement. (A) Cells (yellow) seeded into agarose gels (pink) formed prostate tumor spheroids in the bottom of wells. (B) After 24 hours, spheroids in 6-well plates were surrounded by Superflab bolus (yellow) and irradiated with 6 MV radiation at 5 cm depth in Solid Water (brown). Measurement was performed 24 hours post-irradiation. (C) Clonogenic assay disaggregated spheroids for plating 2D cultures that were fixed and stained after two weeks. (D) AB proliferation assay added AB solution to gels that was extracted for absorbance measurement. (E) FM added solution of green calcein acetoxymethyl and red propidium iodide to gels that fluoresced in live and dead cells, respectively, during enface imaging. (F) Volumetric dOCT imaging acquired forty repeated frames per cross-section. Dynamic signals were extracted from intensity fluctuations via an FFT-based algorithm assigning red (<0.5 Hz) and green (0.5–5 Hz) frequency bands. (G) 3D agent-based simulation of spheroid seeding, irradiation, and measurement applied experimental LQ survival at treatment. SAD: source-to-axis distance; B: Superflab bolus; SW: Solid Water; AB: Alamar Blue; FM: fluorescence microscopy; dOCT: dynamic optical coherence tomography; FFT: Fast-Fourier transform; LQ: linear-quadratic model.

### 2.2 Radiation treatment

A Varian TrueBeam linear accelerator was used to deliver 1, 2, 4, 6, 8, or 10 Gy to monolayer and spheroid cultures with a 6 MV beam 24 hours after seeding (Fig. 1B). Each 6-well plate containing gels with spheroids was positioned at a source-to-axis distance (SAD) of 100 cm, at a depth of 5 cm within Gammex 457-CTG Solid Water, and surrounded by Superflab bolus for scatter conditions. Nonetheless, we expect a delivered dose with an uncertainty in the range of 10%. T-25 culture flasks of monolayer culture were similarly situated and treated. Live control spheroids, which were not exposed to radiation, were removed from incubation and handled identically to treated samples. At the time of treatment, a subset of live control spheroids were fixed with 4% formaldehyde (Thermo Fisher Scientific, USA) in PBS (Wisent Inc., Canada) for 1 hour to eliminate metabolic activity.

These fixed spheroids were subsequently stored at 4°C for approximately 24 hours before measurement to minimize dOCT motion artifacts induced by fixation.

### 2.3 Biological assays

#### 2.3.1 Clonogenic assay

Clonogenic assay was performed with monolayer and spheroid cultures 24 hours post-irradiation to measure clonogenic survival (Fig. 1C). To disaggregate spheroids for clonogenic plating, spheroids were aggressively flushed out of the wells with their surrounding media and transferred into a falcon tube for mechanical vortexing, followed by incubation for two hours to allow cells to settle by gravity. The supernatant was then removed and replaced with at least twice the remaining volume in 0.25% trypsin-EDTA (Thermo Fisher Scientific, USA). After incubation for ten minutes, the trypsin solution was mechanically vortexed again and centrifuged for subsequent cell counting. For monolayer culture, cells were incubated in 0.25% trypsin-EDTA for 3 minutes, then centrifuged for counting. Following cell counting, cells from each experimental condition were plated into 6-well plates at low densities and allowed to grow for approximately two weeks. Colonies were then fixed with 4% formaldehyde in PBS for 1 hour and subsequently stained with 0.5% crystal violet (Thermo Fisher Scientific, USA) dissolved in water for 30 minutes. Stained colonies were counted, and the plating efficiency (PE) was calculated in live control samples as follows:

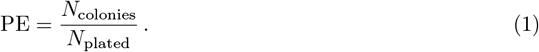

where *N*_colonies_ is the number of colonies counted and *N*_plated_ is the number of cells plated. Then, in treated samples, survival fraction (SF) was calculated as follows:

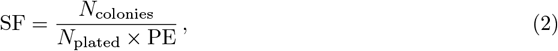

The linear and quadratic parameters *α* and *β* were extracted by least-squares fitting of SF values for each treated dose *d* to the LQ model:

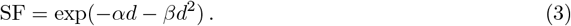

#### 2.3.2 Alamar Blue proliferation assay

AB proliferation assay was performed 24 hours post-irradiation to measure the average cellular proliferation within the tumor spheroids (Fig. 1D). To achieve a final AB concentration of 5% within each 500 *µ*L gel, 34.5 *µ*L of AB (Sigma-Aldrich, USA) and 156 *µ*L of RPMI were added drop-wise to each gel after careful removal of the culture medium in 10 *µ*L increments. Samples were incubated for 3 hours at 37°C with 5% CO_2_ in a humidified atmosphere. Following incubation, the absorbance intensity of the extracted solution was recorded at 570 nm using a BioTek Synergy H1 microplate reader. Three gels containing spheroids were measured for each experimental condition. Gels without cells were processed identically and measured to determine background absorbance. For each measurement of gels containing spheroids, the average background intensity was subtracted, and the resulting AB absorbance was normalized relative to the absorbance measured in the live control samples.

### 2.4 Fluorescence microscopy

FM was performed 24 hours post-irradiation to visualize cell death within spheroids using the Live/Dead Cell Viability assay (Sigma-Aldrich, USA). Live cells were labeled using calcein acetoxymethyl (calcein-AM), which fluoresces green through intracellular processes [48]. Dead cells were labeled using propidium iodide (PI), which fluoresces red upon binding with DNA or RNA that is inaccessible in live cells [49]. After careful removal of culture medium, a solution containing 0.2875 *µ*L calcein-AM, 1.15 *µ*L PI, 95 *µ*L RPMI, and 95 *µ*L PBS was added to each gel. Samples were then incubated for one hour at 37°C with 5% CO_2_ in a humidified atmosphere. Following incubation, gels were transferred to coverslip-bottomed Petri dishes and imaged using a Zeiss Axio Observer widefield microscope that was operated by ZEN2 Blue Edition software and equipped with an Axiocam 506 mono camera. A Plan-Apochromat 63/1.4 Oil Ph3 M27 objective (Carl Zeiss Microscopy LLC, USA) was used to acquire fluorescent images, while an incubation plate (PECON, Germany) maintained the temperature at 37°C during imaging.

### 2.5 OCT

Tumor spheroids were imaged with the LF-dOCT system 24 hours post-irradiation (Fig. 1F). A detailed description of the system design and performance can be found in [47]. Briefly, the system employed a broadband superluminescent diode (SLD) light source (cBLMD-T-850-HP, Superlum, Ireland) with a central wavelength of 842 nm and a full-width-at-half-maximum (FWHM) spectral bandwidth of 179 nm, achieving an axial resolution of 2.6 *µ*m in air. A 5× microscope objective (M Plan APO, Mitutoyo, Japan) provided a lateral resolution of 6.4 *µ*m. The system operated at an image acquisition speed of 2,000 frames per second (fps), and had approximately 93 dB sensitivity, measured with an incident power of 3.5 mW at the image plane.

Volumetric dOCT images were acquired and processed following previously described methods [50]. Each dOCT volume consisted of 400 B-scans, with each B-scan composed of 400 A-scans. The dOCT volume acquisition was divided into four sub-volumes, each imaged with 40 repetitions at a repetition rate of 10.7 Hz. OCT images of 15 tumor spheroids per experimental condition were acquired, with each spheroid imaged for approximately 16 seconds during dOCT recording. Spheroids were maintained at 21°C without CO_2_ supply during imaging. All OCT and dOCT images presented for each experimental condition were generated from the same representative spheroid, whereas different spheroids were used for FM.

Morphological OCT images were generated using a standard OCT image reconstruction process and are displayed in logarithmic scale with the surrounding gel masked. Volumetric OCT masks were analyzed numerically to estimate the volume *V*_sph_ and external surface area *S* of each spheroid, as well as the volume of internal gaps *V*_gap_ enclosed by that surface area. Sphericity, which reaches a maximum value of 1 for a perfect sphere, was calculated to compare the external surface area of each spheroid to the volume it enclosed as follows:

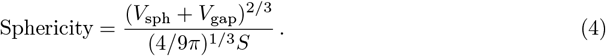

To generate dOCT images, a Hann window function was first applied along the time direction to the linearly scaled OCT intensity values at each pixel location. Following zero-padding of the data, a Fast Fourier Transform (FFT) was performed along the time-axis [51, 52]. Next, integrals were calculated over two distinct frequency bands: a lower-frequency (“slow”) red band below 0.5 Hz and a higher-frequency (“fast”) green band between 0.5 and 5 Hz. After applying logarithmic scaling to each of the red and green dOCT channels, a small region of culture medium located above each spheroid in a central B-scan was selected. The sum of the mean and standard deviation of the intensity within this reference region was subtracted from each pixel location. To quantitatively analyze dynamic activity, a raw dynamic signal metric was computed for each masked spheroid as follows:

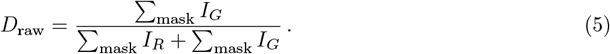

Here, *I*_*R*_ and *I*_*G*_ represent the intensity values of the red and green channels, respectively. The average raw dynamic signal measured in the formaldehyde-fixed spheroids was subtracted from the corresponding signal measured in live spheroids, and the resulting value was normalized relative to live control as follows:

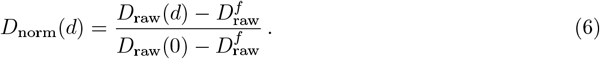

Here, *D*_raw_(*d*) and 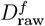 represent the raw dynamic signals measured in spheroids treated with radiation dose *d* (Gy) and formaldehyde-fixed spheroids, respectively. Due to attenuation with depth caused by spheroid density, the numerical analysis excluded any enface (lateral view) planes whose average intensity was lower than the median intensity across the entire volume. To display dOCT images, the masked red and green channels were independently normalized to those of a representative spheroid before a Gaussian FFT filter was applied with *σ*_*R*_ = 90 and *σ*_*G*_ = 65 respectively. Finally, a 3 × 3 median filter was applied to each channel before combining them into an RGB color image.

### 2.6 Simulation

For this study, we also developed an agent-based computer model of 3D tumor spheroid growth and radiation treatment using the software CompuCell3D [53] (Fig. 1G). Simulated time evolved according to a Monte-Carlo method, which minimized the effective energy of the 3D lattice containing the spheroid. Each simulation began with a suspension of 1,000 well oxygenated *Normoxic* tumor cells placed randomly inside the gel within a curved well described by:

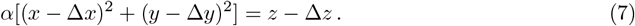

Here, as extracted from OCT images, *α* = 0.0025 *µ*m^−1^, and (Δ*x*, Δ*y*, Δ*z*) positioned the bottom of the well at the center of the bottom surface inside a simulation space measuring 810 *µ*m across and 405 *µ*m deep with 5 *µ*m voxels. Initially suspended cells settle under gravity and self-assemble into a spheroid, driven by the energetic preference of live cells to be in contact with other live cells rather than the surrounding gel or culture medium. Oxygen, governed by a PDE, was maintained at a constant partial pressure in the culture medium, and diffused through the spheroid with a diffusion coefficient approximately equivalent to half that of oxygen in water. Simulated cells consumed oxygen, and their growth rate depended on the local partial pressure of oxygen according to a Michaelis-Menten saturation curve. When a cell grew to a sufficient volume, it divided into two daughter cells of equal volume. Due to competition and limited oxygen diffusion within the spheroid bulk, *Normoxic* cells reversibly became *Hypoxic* when oxygen-deprived, and irreversibly became *Necrotic* upon severe oxygen deficiency. The simulation time scale was calibrated so that *Normoxic* cells divided approximately every 24 hours, and spheroid dimensions were comparable to those measured by longitudinal OCT imaging 24 hours after seeding [50].

Radiation treatment was simulated 24 hours after seeding. At the treatment time, each cell within the spheroid probabilistically survived according to the LQ curve derived from spheroid clonogenic experiments. Cells killed by radiation irreversibly became *Apoptotic* only upon attempted cell division. The simulation explicitly distinguished cell death due to necrosis by nutrient deprivation from radiation-induced apoptosis. However, *Necrotic* and *Apoptotic* cells behaved identically and, unlike *Normoxic* and *Hypoxic* cells, did not preferentially adhere to neighboring cells. Formaldehyde fixation was not explicitly simulated; instead, it was approximated by assuming all cells in the live control simulation became *Apoptotic* at the time of treatment.

Numerical morphological and dynamic analysis of simulated live control spheroids and spheroids irradiated with 1, 2, 6, and 10 Gy were performed 24 hours post-irradiation. Morphological metrics including spheroid volume, surface area, and internal gap volume were calculated voxel-wise using the 3D lattice and were subsequently used to compute sphericity according to Eq. (4). Simulated SF was calculated using Eq. (2) by estimating *N*_colonies_ and *N*_plated_ as the number of living cells and the total number of cells in the spheroid, respectively. Due to the stochastic nature of the Monte Carlo simulation, each condition was simulated ten times. A detailed mathematical description of the simulation can be found in the supplementary data.

### 2.7 Statistics

All measurements are presented as averages with corresponding error bars. Clonogenic SF error was estimated by Fieller’s theorem with a 95% confidence interval from a Poisson distribution, as described by Gupta et al. [54]. For comparison with alternative SF measurements, clonogenic SF was presented using the LQ fit. The upper (lower) LQ error bounds were calculated using LQ parameters, *α* and *β*, that were fit to experimental clonogenic SF values plus (minus) the SF error estimated by Fieller’s theorem. All other error bars represent standard deviation.

Numerical OCT and dOCT measurements that fell outside of the interquartile range were excluded from the analysis. In the numerical morphological analysis, one-way ANOVA tests were performed to compare live control spheroids with spheroids irradiated with 10 Gy and those fixed with formaldehyde for each metric. If the omnibus ANOVA test was significant, independent Welch’s t-tests were conducted to compare each tested condition individually against the live control spheroids. In the numerical dynamic analysis, independent Welch’s t-tests were performed to compare the normalized SF measured by dOCT and AB proliferation assay for each dose of radiation. A significance level *α* of 0.05 was employed and t-tests were Bonferroni-corrected.

Statistically significant t-test results are indicated on the relevant plots with an asterisk.

## 3 Results

Figure 2 summarizes the volumetric OCT and simulated morphological analyses of live control, irradiated, and formaldehyde-fixed spheroids. The first and second rows of Figure 2 present the average spheroid volume, internal gap volume, external surface area, and sphericity calculated voxel-wise in masked volumetric OCT images and simulation. As the radiation dose increased from 0 Gy to 6 Gy, simulations predicted reductions in spheroid volume, internal gap volume, and surface area, alongside an increase in sphericity (Fig. 2A-D). All simulated morphological metrics then plateaued between 6 Gy and 10 Gy, with 10 Gy significantly different from live control (Fig. 2A-D). Compared to the OCT measurements, simulated gap volumes (Fig. 2B) were three orders of magnitude larger, while surface area (Fig. 2C) and sphericity (Fig. 2D) values were one order of magnitude larger and smaller, respectively. Despite these differences in magnitude, the general trends of the morphological metrics observed by volumetric OCT were consistent with simulated results for radiation doses between 0 Gy and 2 Gy (Fig. 2A-D). However, in the OCT analysis, spheroid volume, gap volume and surface area increased at 6 Gy, then plateaued at 10 Gy, and further increased in formaldehyde-fixed spheroids (Fig. 2A-C). Notably, the surface area of formaldehyde-fixed spheroids was significantly larger than that of live control spheroids (Fig. 2C). In contrast, after minimal changes between 2 Gy and 6 Gy, both simulated and OCT-measured sphericity achieved a maximum at 10 Gy and minimum in formaldehyde-fixed spheroids; both extremes were significantly different from the live control spheroids (Fig. 2D).

**Figure 2.**
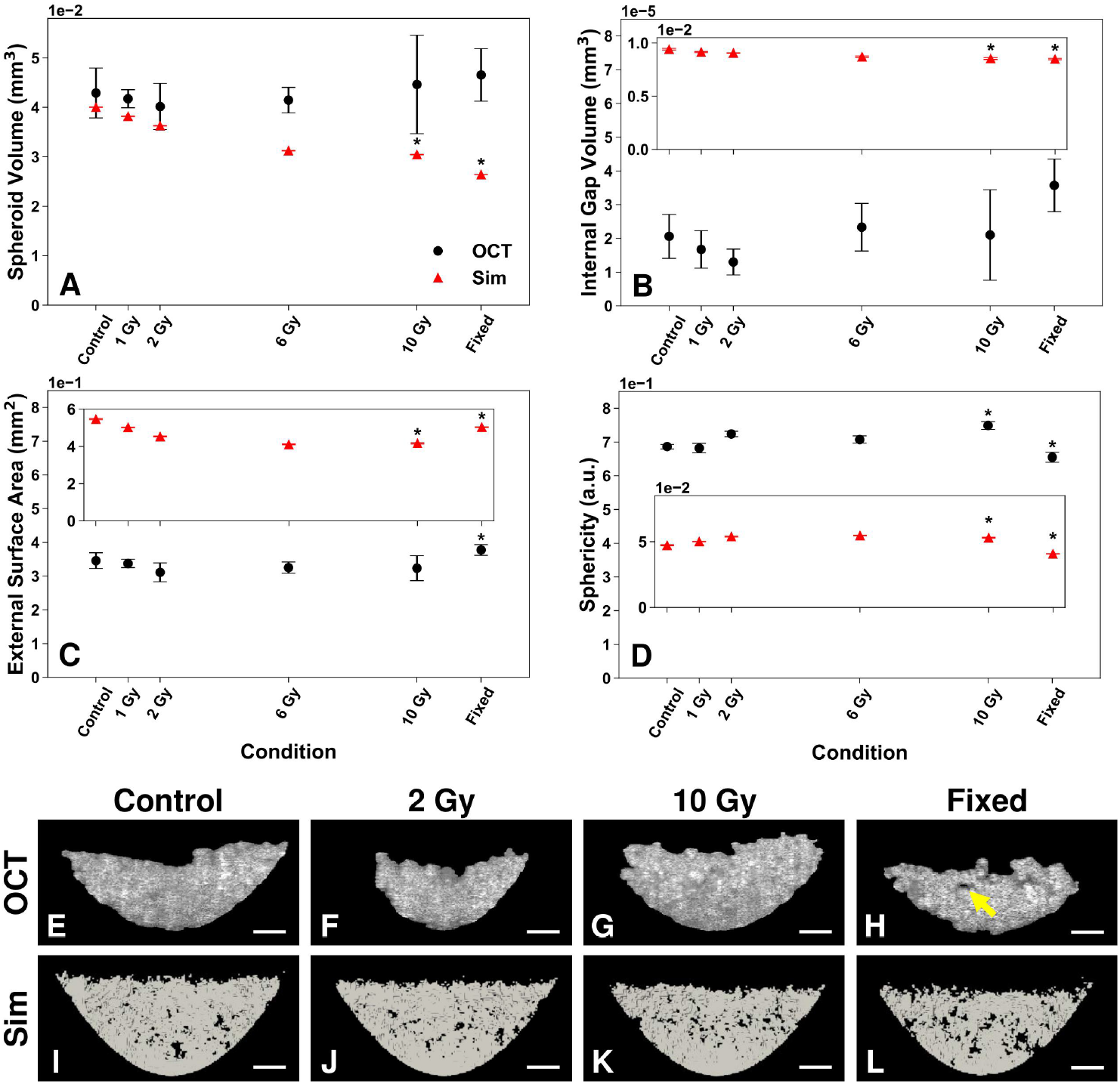
Morphological OCT and simulated analyses of live control, irradiated (1, 2, 6, and 10 Gy), and formaldehyde-fixed prostate tumor spheroids. Numerical analysis of (A) spheroid volume, (B) internal gap volume, (C) external surface area, and (D) sphericity given by Eq. (4). Average OCT-measured (black circle) and simulated (red triangle) values are presented with standard deviation error bar. Simulated data in (B-D) are inset. Asterisks indicate significant differences (10 Gy or fixed vs. live control; Methods Section 2.7). Central (third row) masked OCT B-scans and (fourth row) simulated visualizations of live control, irradiated (2 and 10 Gy), and formaldehyde-fixed spheroids. An internal gap is labeled with yellow arrow. Sim: simulation. Scale bars: 100 *µ*m.

The third and fourth rows of Figure 2 show central cross-sectional masked OCT images and corresponding simulated visualizations, respectively. OCT images exemplify the smaller volume and surface area of spheroids treated with 2 Gy (Fig. 2F) compared to 10 Gy (Fig. 2G), which is of a comparable size to live control (Fig. 2E). An internal gap can be observed in the OCT image of a formaldehyde-fixed spheroid (Fig. 2H, yellow arrow), which otherwise resembles the live control and irradiated spheroids. In contrast, simulated spheroids display largely uniform cross-sectional shapes and sizes across all radiation doses (Fig. 2I-K). However, simulated formaldehyde-fixed spheroids similarly contain more internal gaps than both control and irradiated spheroids (Fig. 2L).

Figure 3 summarizes the dynamic analysis. Clonogenic SF of prostate cancer cells cultured in 3D spheroids and conventional 2D monolayers are presented in Figure 3A. Compared to 2D monolayer culture, cells cultured in spheroids demonstrated higher radioresistance with approximately 6% and 2% clonogenic survival after irradiation with 6 Gy and 10 Gy, respectively (Fig. 3A). Figure 3B compares the SF of spheroids measured via normalized dynamic signals derived from masked volumetric dOCT images, normalized absorbance from AB proliferation assay, simulations, and clonogenic assay. Generally, SF values measured by dOCT were larger than those measured by AB but the differences were not statistically significant; initially increasing slightly with radiation dose from live control, SF values measured by both methods then decreased and subsequently reached a maximum at 10 Gy (Fig. 3B). In contrast, clonogenic and simulated SF values consistently decreased with increasing dose, with clonogenic SF decreasing at a faster rate (Fig. 3B). The second and third rows of Figure 3 show volumetric dOCT images and simulated spheroids irradiated with 1, 2, 6, and 10 Gy, respectively. In dOCT volumetric images, high frequency (green channel) intensity was predominantly observed throughout the spheroid bulk across all radiation doses, reaching its maximum intensity in the spheroid treated with 10 Gy (Fig. 3C-F). Conversely, simulated volumetric images demonstrated a distinct hypoxic core in the live control spheroids (Fig. 3G), with an increasing proportion of *Apoptotic cells* distributed throughout the spheroid bulk as radiation dose increased (Fig. 3H-J).

**Figure 3.**
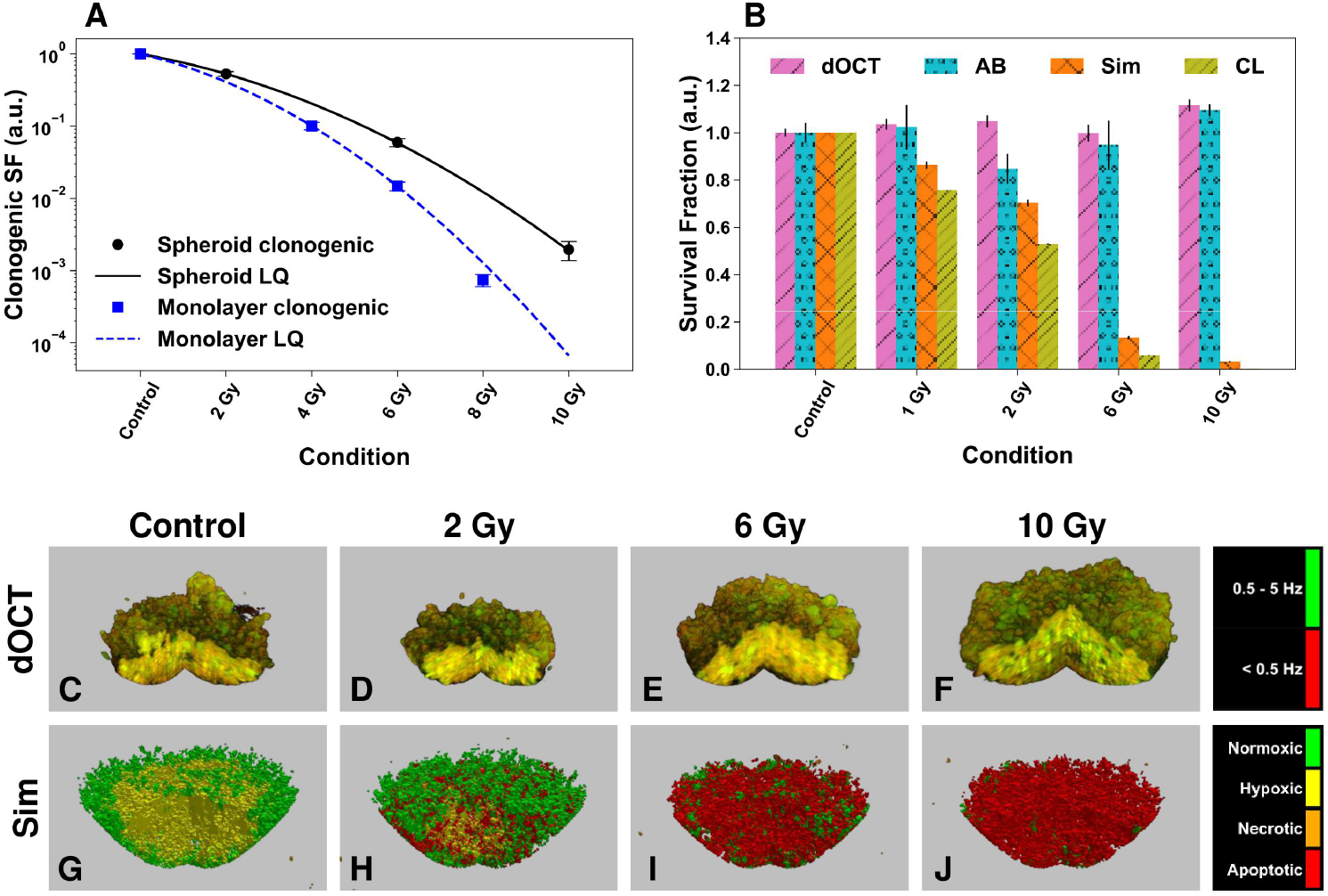
Dynamic analysis of live control and irradiated prostate tumor spheroids. (A) Clonogenic assay SF values with LQ model fit line for 3D spheroids (black circle, solid line) and 2D monolayer culture (blue square, dashed line). (B) SF of spheroids measured via normalized dynamic signal from volumetric dOCT images (purple /hatch), normalized absorbance from AB proliferation assay (blue circle hatch), simulated clonogenic assay S_*CL*_ (orange X hatch), and LQ fit from clonogenic assay (olive //hatch). Data shown are averages with error bars as described in Methods Section 2.7. Volumetric dOCT images (second row) and simulated visualizations (third row) of live control and irradiated (2, 6, and 10 Gy) spheroids. A central slice has been cut out of the volumes. Colormaps for dOCT (red: <0.5 Hz, green: 0.5-5 Hz); and simulation (red: *Apoptotic*, orange: *Necrotic*, yellow: *Hypoxic*, green: *Normoxic* cells). SF: survival fraction; LQ: linear-quadratic model; AB: Alamar Blue; Sim: simulation; CL: clonogenic.

Figure 4 compares OCT, dOCT, FM, and simulated enface images of live control, irradiated, and formaldehyde-fixed spheroids. OCT, dOCT, and FM enface images showed minimal variation between live control spheroids and those treated with radiation; however, live control and irradiated spheroids appeared significantly different from formaldehyde-fixed spheroids (Fig. 4). Individual cells could be resolved in OCT enface images, with live control and irradiated spheroids appearing generally compact (Fig. 4A-E), whereas formaldehyde-fixed spheroids displayed noticeable internal gaps (Fig. 4F, yellow arrow). Similarly, individual cells and gaps were clearly resolved in dOCT images (Fig. 4G-L, yellow arrow). In dOCT images, green channel intensity was higher at the spheroid periphery and increased slightly with increasing radiation dose in live control and irradiated spheroids (Fig. 4G-K). In contrast, formaldehyde-fixed spheroids exhibited ubiquitous red channel intensity with only a few individual cells appearing green (Fig. 4L). In FM enface images, the live control and irradiated spheroids appeared predominantly green, with some central red fluorescence (Fig. 4M-Q). However, after formaldehyde fixation, the FM image was dominated by red fluorescence and exhibited minimal central green fluorescence (Fig. 4R). Unlike OCT, dOCT, and FM images, simulated spheroids were comprised of a progressively increasing proportion of red *Apoptotic* cells as radiation dose increased (Fig. 4S-W) and appeared entirely red following simulated fixation (Fig. 4X).

**Figure 4.**
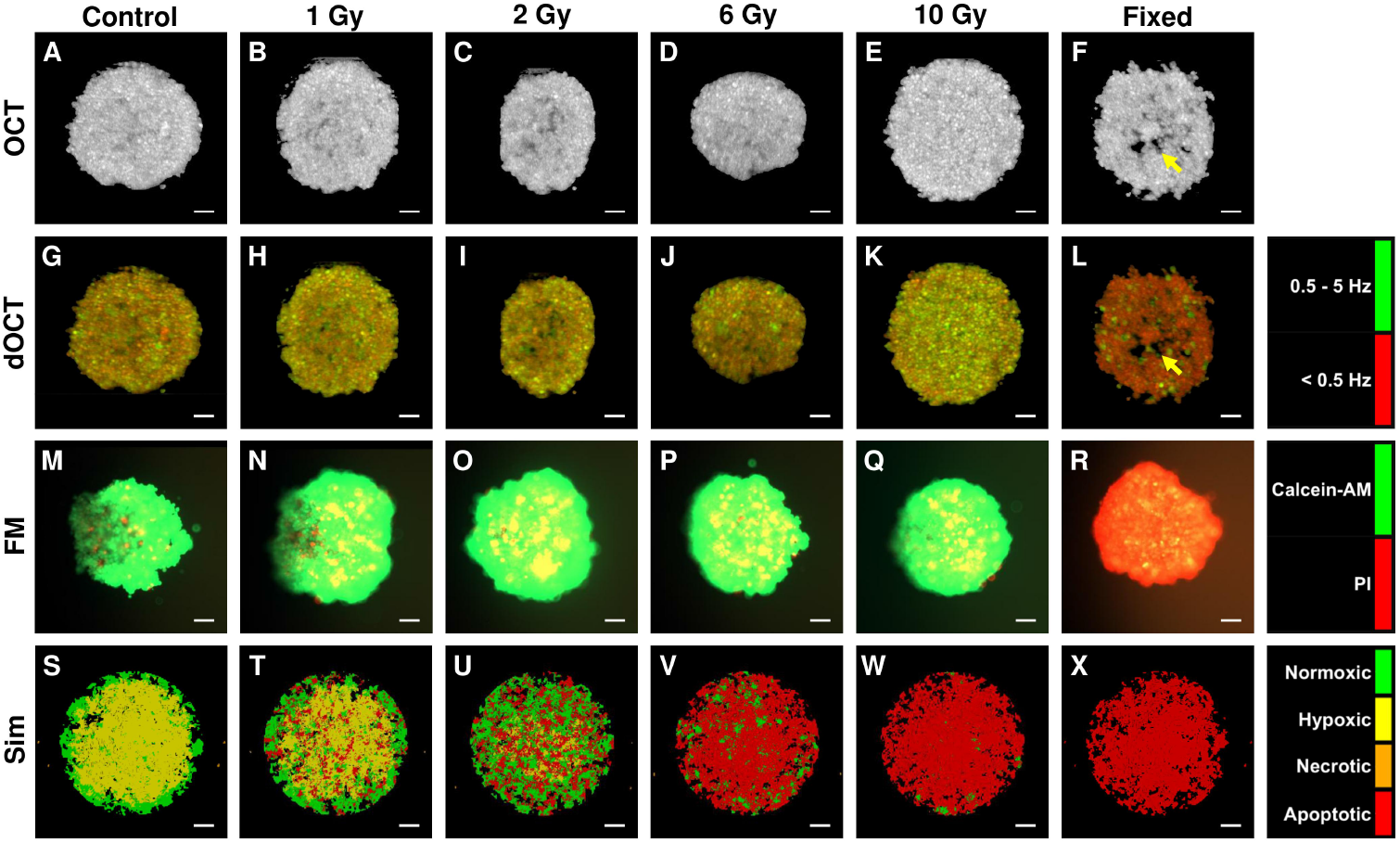
OCT maximum projection (first row), dOCT maximum projection (second row), FM (third row), and simulated (fourth row) enface images of live control, irradiated (1, 2, 6, and 10 Gy), and formaldehyde-fixed prostate tumor spheroids. An internal gap is labeled with yellow arrow. Colormaps for dOCT (red: <0.5 Hz, green: 0.5-5 Hz); FM (red: PI, green: calcein-AM); and simulation (red: *Apoptotic*, orange: *Necrotic*, yellow: *Hypoxic*, green: *Normoxic*). FM: fluorescence microscopy; propidium iodide: PI; calcein-AM: calcein acetoxymethyl; Sim: simulation. Scale bars: 100 *µ*m.

## 4 Discussion

### 4.1 Morphological measurement of radiation effects

Simulations provide a platform for visualizing and precisely quantifying the outcomes of logical experiments conducted under perfectly understood, albeit oversimplified, biological conditions [26]. Our 3D agent-based simulation assumed the spheroid grew within a symmetric paraboloid-shaped well given by Eq. (7) embedded in an agarose gel, with uniform oxygen diffusion and uniform oxygen consumption per unit volume of living cells. At the time of simulated radiation treatment, each cell survived irradiation with a probability equal to the SF given by the experimentally derived spheroid LQ curve shown in Figure 3A. Under these assumptions, we observed that simulated spheroid volume (Fig. 2A), internal gap volume (Fig. 2B), and external surface area (Fig. 2C) decreased with increasing radiation dose, subsequently plateauing between 6 Gy and 10 Gy. According to the LQ model, the probability of cell death increased exponentially with increasing dose (Fig. 3A), resulting in a corresponding increase in the number of simulated *Apoptotic* cells at 24 hours after treatment (Fig. 4S-W). Simulated *Apoptotic* cells failed to undergo cell division; thus, spheroids treated with lower doses exhibited larger volume and surface area due to a greater number of successful cell divisions occurring between treatment and analysis. The reduced internal gap volume in simulated spheroids treated with higher doses resulted from structural collapse caused by an increased proportion of non-adherent *Apoptotic* cells. Simulated sphericity increased with dose due to its higher sensitivity to surface area per Eq. (4), which decreased faster than the combined spheroid and internal gap volumes (Fig. 2A-D). Despite a 10% decrease in simulated SF between 6 Gy and 10 Gy (Fig. 3B), the corresponding morphological changes were negligible (Fig. 2C-D) because the simulated spheroids irradiated with 6 Gy were already predominantly composed of *Apoptotic* cells (Fig. 4V).

Simulated surface areas and internal gap volumes were significantly larger than those measured with volumetric OCT masks (Fig. 2B-C). Simulations precisely quantified these metrics with a voxel resolution of 5 *µ*m, whereas noise and an imperfect masking process complicated interpretation of the OCT measurements, despite our system demonstrating similar axial and lateral resolutions of 2.6 *µ*m and 6.4 *µ*m, respectively. The observed discrepancy in surface area illustrates the coastline paradox, wherein using higher resolutions to measure fractal-like surfaces, such as coastlines or spheroid surfaces, paradoxically increases the measured value. Rather than improving accuracy, the higher resolution of the simulation merely accounts for increasingly smaller surface features that the OCT mask smooths out. Furthermore, because OCT masks prioritize containing all cells within the spheroid, the smoothing process preferentially reduces internal gap volume and consequently overestimates spheroid volume. Underestimation of gap volume by OCT masking resulted in greater discrepancy with simulation compared to its overestimation of spheroid volume since the volume affected represents a substantially larger fraction of total gap volume than of total spheroid volume (Fig. 2A-B).

Despite these differences in absolute magnitude, morphological trends observed in simulated spheroids irradiated with low doses corresponded well with OCT analysis. Raitanen et al. [21] similarly observed no significant size differences in PC3 spheroids irradiated with doses up to 8 Gy after 3 days. However, morphological discrepancies between simulation and OCT increased with increasing dose. Agreement was strongest for live control spheroids because the simulation was calibrated to spheroid dimensions measured in a longitudinal OCT study of untreated spheroids [50]. With higher doses, the simplified mechanism of cell death and unrealistically instantaneous apoptotic process in the simulation diverged increasingly from the complex biological processes observed experimentally.

### 4.2 Clonogenic measurement of radiation effects

In reality, radiation-induced cell death is neither instantaneous nor straightforward. Radiation damages DNA indirectly via reactive oxygen species (ROS) and directly via ionizing tracks that create single-strand breaks (SSBs) and more lethal double-strand breaks (DSBs) [8, 55, 56]. DNA damage triggers various signaling pathways that instigate DNA repair [57, 58], the duration of which increases with the complexity of the damage and may extend beyond 24 hours [59, 60]. Radiation exposure also induces G2/M phase arrest, delaying the onset of mitosis by prolonging the G2 phase of the cell cycle, thus preventing cell division with damaged DNA [61, 62]. Persistent cell cycle arrest leads to senescence, a state in which cells stop proliferating but can remain metabolically active [63]. Cells with irreparable DNA damage may undergo mitotic catastrophe and subsequently die through some combination of apoptosis and necrosis [8, 56, 57, 63]. Thus, radiation results in clonogenic death by preventing cells from successfully performing cell division, although cells can be observed to remain metabolically active for some time after treatment [21, 33].

Consequently, the clonogenic assay remains the gold standard method for measuring cell survival after irradiation, as it directly assesses the clonogenic potential of individual cells. We measured higher clonogenic SF in prostate cancer cells cultured in 3D spheroids compared to conventional 2D monolayer culture (Fig. 3A). The improved clonogenic survival of spheroids compared to monolayer culture was first reported by Durand and Sutherland [19] who attributed this phenomenon to increased cell-to-cell contact and it has since been documented across various cell lines [20, 21, 33, 64–67]. Additionally, irradiated spheroids exhibit reduced DNA damage [21, 66] and apoptosis [67] compared to monolayer culture. It has been suggested that the increased survival observed in spheroids may result from the presence of a quiescent hypoxic core, which has been widely documented [20, 22, 23, 33, 68]. Clinically, hypoxia significantly contributes to tumor radio-resistance, in part due to the radio-sensitizing effect of oxygen [69–71]. Cells have been observed to become quiescent under stress such as hypoxia [23, 72], and quiescence itself is known to confer radio-resistance [20, 23]. However, PC3 cells form spheroids as loose aggregates that are not tightly adherent [73, 74], and our FM results showed only a small necrotic core within live control spheroids (Fig. 4M). Therefore, compared to other cells lines, the contribution of a quiescent hypoxic core to the increased radio-resistance observed in PC3 spheroids relative to monolayer culture might be relatively minor.

### 4.3 Alternative measurement of radiation effects

A rapid, simple, and reliable alternative to the clonogenic assay for measuring radiation survival has long been sought [45]. The search for alternative methods is particularly salient for 3D spheroid culture, given the observation by Onozato et al. [20] that the spheroid disaggregation required for clonogenic assay can itself influence radiation response. Our study explored dOCT as another potential alternative alongside proliferation assay for assessing radiation effects in intact spheroids.

We observed that the normalized SF values measured by dOCT were consistently higher than AB proliferation assay at each radiation dose, although these differences did not reach statistical significance (Fig. 3B). The dOCT method effectively distinguished intensity fluctuations due to cellular metabolism, clearly differentiating between control (Fig. 4G) and formaldehyde-fixed spheroids (Fig. 4L). Similarly, AB reduction is a well-established indicator of cellular metabolism [31]. However, the SF values measured by both dOCT and AB were substantially higher than those determined by clonogenic assay (Fig. 3B). While dOCT imaging and AB proliferation assay are more straightforward and efficient to perform than clonogenic assay, they capture cellular activity at only a single time point, which limits their detection of delayed radiation effects [21, 33, 57]. Indeed, the numerous signaling pathways activated by radiation repair [57, 58] and subsequent cell death [63, 75, 76] may increase cellular activity observed with dOCT and AB in the first 24 hours following irradiation. Furthermore, radiation has been reported to activate invasive and proliferative pathways mediated through epidermal growth factor receptor (EGFR) [77], integrins [78], and hypoxia inducible factor (HIF) [79], potentially increasing measured cellular activity. Moreover, dOCT measurements are sensitive not only to metabolic activity but also to physical changes in cell morphology and volume. Apoptotic cells undergo characteristic physical transformations, including shrinking, surface blebbing, and chromatin condensation (pyknosis), before nucleus fragmentation (karyorrhexis) [75]. In contrast, necrotic cells exhibit cell swelling, undergo nuclear disintegration (karyolysis), and ultimately degrade, releasing their contents [63, 75]. These physical transformations could explain the higher SF measured by dOCT compared to AB; however, the investigation of post-radiation physical changes such as blebbing and pyknosis would benefit from higher-resolution OCT imaging focused on specific regions rather than the entire spheroid volume. Additionally, high-frequency motion detected by dOCT could also reflect motion artifacts unrelated to cellular behavior, such as Brownian motion, or external scanner and camera noise.

To accurately estimate clonogenic survival with proliferation assay, Nikzad and Hashemi [10] recommended continuous measurement for several days following irradiation to characterize the radiation-induced growth delay. Future work should similarly include multiple dOCT measurements in the week following treatment to assess the delayed effect of radiation. Such growth-delay characterization could then be directly compared with clonogenic SF measurements, analogous to the approach by Buch et al. [30] using MTT proliferation assay.

The simulated SF values were lower than those measured by dOCT and AB proliferation assay, despite all methods estimating instantaneous survival at a single time point 24 hours post-irradiation (Fig. 3B). The discrepancy is likely due to the simplified model of radiation-induced cell death. In our model, hypoxia indirectly delayed radiation effects by slowing cellular growth, delaying attempted cell division, and consequently delaying apoptosis. However, during simulated irradiation, *Hypoxic* cells did not exhibit radio-resistance. In contrast, other models have explicitly included oxygen effects on radiation survival by modifying LQ survival with an oxygen saturation curve [80, 81]. Various investigators have also explicitly modeled the process of DNA damage repair [81–83], growth delay and cell cycle arrest [81, 84], as well as the effect of cell cycle phase on radiation survival [85–89]. In this regard, agent-based models provide a particularly effective platform to characterize tumor heterogeneity and explore its implications in radiation treatment.

The clonogenic assay inherently integrates these complex biological factors into a binary probability of long-term clonogenic survival, which we simulated probabilistically at the time of treatment. However, simulated SF remained higher than clonogenic SF for all doses (Fig. 3B) because not all cells killed by radiation had attempted cell division and become *Apoptotic* by the measurement time point, 24 hours post-irradiation. Additionally, our model assumed instantaneous cell death without explicitly modeling the duration of apoptosis, a process estimated to take approximately 24 hours [76]. At the measurement time point, many cells classified as *Apoptotic* in the simulation would realistically still be undergoing the physically and metabolically dynamic process of cell death, potentially contributing to the elevated SF values observed by both dOCT and AB proliferation assay.

### 4.4 Comparison of causes of cell death

Our study investigated multiple causes of cell death. At the time of radiation treatment, live control spheroids were also fixed with 4% formaldehyde to eliminate metabolic activity. Clonogenic assay demonstrated approximately 2% clonogenic survival in spheroids irradiated with 10 Gy (Fig. 3B), and FM revealed widespread cell death in formaldehyde-fixed spheroids (Fig. 4R). However, dOCT and FM images of control and irradiated spheroids both showed higher green channel intensity along the spheroid periphery with some central red channel intensity, whereas formaldehyde-fixed spheroids appeared almost entirely red in color (Fig. 4G-R). Both imaging techniques were applied 24 hours after treatment, at a time when irradiated cells remained metabolically active and structurally intact [21, 33, 57]. In contrast, fixation-induced cell death occurred quickly and efficiently. Thus, dOCT accurately detected the metabolically inactive, fixed spheroids as dynamically static, consistent with the extensive cell death observed by FM.

Morphologically, formaldehyde fixation preserved spheroid structure without significant tissue shrinkage by cross-linking proteins [90], effectively arresting spheroid morphology at the treatment time. Due to the simplified nature of our simulation, fixation was not directly simulated; instead, we approximated fixation by assuming all cells in live control spheroids at the treatment time were *Apoptotic* (Fig. 4X). At the time of treatment and fixation, 24 hours after seeding, the spheroids were still condensing to become more tightly aggregated [50]. This process is reflected in the larger OCT-measured spheroid volume (Fig. 2A), internal gap volume (Fig. 2B), and external surface area (Fig. 2C) of formaldehyde-fixed spheroids compared to live control, although the difference was only statistically significant for surface area. In contrast, the simulated formaldehyde-fixed spheroids had significantly smaller volume (Fig. 2A), internal gap volume (Fig. 2B), and external surface area (Fig. 2C) compared to live control spheroids. The morphological difference between OCT measurement and simulation is likely due to the simplified modeling of the complicated and prolonged process of spheroid aggregation. Consequently, simulated spheroids were smaller and more tightly aggregated at the time of treatment and fixation, and they remained so in the complete absence of simulated cell division and growth in the 24-hour interval between fixation and analysis.

## 5 Conclusion

In conclusion, we employed a spectral-domain LF-dOCT system to acquire volumetric dOCT images of prostate tumor spheroids 24 hours following radiation treatment. Morphological trends measured by volumetric OCT analysis aligned with simulations of a novel 3D agent-based model of radiation treatment whose parameters were informed by experimental clonogenic assay data. The effects of radiation measured using dOCT showed excellent quantitative and qualitative agreement with conventional biological techniques, AB proliferation assay and FM, respectively. The strong agreement between dOCT and AB proliferation assay suggests that a longer duration of repeated dOCT measurement in tumor spheroids following radiation treatment could offer a non-invasive alternative to the clonogenic assay, eliminating the need for spheroid disaggregation.

## Supporting information

Supplemental Model Information

## Acknowledgments

It is with sadness that the authors acknowledge the recent passing of the senior author, Dr. Kostadinka Bizheva. The authors would like to gratefully acknowledge Dr. Mohammad Kohandel for generous use of his laboratory equipment. We thank Dr. Brian Ingalls and Atiyeh Ahmadi for their expertise and use of their fluorescent microscope, and Dr. Qing-Bin Lu and Olya Changizi for access and assistance with their microplate reader. Lastly, we are grateful for the help of Catherine McKenna for assisting in cell culture preparation.

## Data availability

Data underlying the results presented in this paper are not publicly available at this time but may be obtained from the authors upon reasonable request.

## Code availability

The code used to process the OCT and dOCT images is publicly available on GitHub at github.com/skswanso/D-OCT.git. The model code used for simulation is also publicly available on GitHub at github.com/skswanso/CPM.git.

## References

1. Siegel RL, Kratzer TB, Giaquinto AN, Sung H, Jemal A. Cancer statistics, 2025. Ca. 2025;75(1):10.

2. Brenner DR, Gillis J, Demers AA, Ellison LF, Billette JM, Zhang SX, et al. Projected estimates of cancer in Canada in 2024. CMAJ. 2024 May;196(18):E615–23. Available from: https://www.cmaj.ca/content/196/18/E615.

3. Keyes M, Crook J, Morton G, Vigneault E, Usmani N, Morris WJ. Treatment options for localized prostate cancer. Canadian Family Physician. 2013;59(12):1269–74.

4. Withers HR. The four R’s of radiotherapy. In: Advances in radiation biology. vol. 5. Elsevier; 1975. p. 241–71.

5. Steel GG, McMillan TJ, Peacock J. The 5Rs of radiobiology. International journal of radiation biology. 1989;56(6):1045–8.

6. Brown JM, Carlson DJ, Brenner DJ. The tumor radiobiology of SRS and SBRT: are more than the 5 Rs involved? International Journal of Radiation Oncology* Biology* Physics. 2014;88(2):254–62.

7. Brenner DJ, Sachs RK, Peters LJ, Withers HR, Hall EJ. We forget at our peril the lessons built into the α/β model. International journal of radiation oncology, biology, physics. 2012;82(4):1312–4.

8. Hall E. Radiobiology for the radiologist. Lippincott Williams & Wilkins; 2006.

9. Puck TT, Marcus PI. Action of x-rays on mammalian cells. The Journal of experimental medicine. 1956;103(5):653–66.

10. Nikzad S, Hashemi B. MTT assay instead of the clonogenic assay in measuring the response of cells to ionizing radiation. J Radiobiol. 2014;1(1):3–8.

11. K Sachs PH, DJ Brenner R. Review the link between low-LET dose-response relations and the underlying kinetics of damage production/repair/misrepair. International journal of radiation biology. 1997;72(4):351–74.

12. Brenner DJ, Hlatky L, Hahnfeldt P, Huang Y, Sachs R. The linear-quadratic model and most other common radiobiological models result in similar predictions of time-dose relationships. Radiation research. 1998;150(1):83–91.

13. Carlson DJ, Stewart RD, Semenenko VA, Sandison GA. Combined use of Monte Carlo DNA damage simulations and deterministic repair models to examine putative mechanisms of cell killing. Radiation research. 2008;169(4):447–59.

14. Pampaloni F, Reynaud EG, Stelzer EH. The third dimension bridges the gap between cell culture and live tissue. Nature reviews Molecular cell biology. 2007;8(10):839–45.

15. Weiswald LB, Bellet D, Dangles-Marie V. Spherical cancer models in tumor biology. Neoplasia. 2015;17(1):1–15.

16. Nath S, Devi GR. Three-dimensional culture systems in cancer research: Focus on tumor spheroid model. Pharmacology & Therapeutics. 2016 Jul;163:94–108. Available from: https://www.sciencedirect.com/science/article/pii/S0163725816300213.

17. Baker BM, Chen CS. Deconstructing the third dimension – how 3D culture microenvironments alter cellular cues. Journal of Cell Science. 2012 Jul;125(13):3015–24. Available from: 10.1242/jcs.079509.

18. Costa EC, Moreira AF, de Melo-Diogo D, Gaspar VM, Carvalho MP, Correia IJ. 3D tumor spheroids: an overview on the tools and techniques used for their analysis. Biotechnology Advances. 2016 Dec;34(8):1427–41. Available from: https://www.sciencedirect.com/science/article/pii/S0734975016301379.

19. Durand R, Sutherland R. Effects of intercellular contact on repair of radiation damage. Experimental cell research. 1972;71(1):75–80.

20. Onozato Y, Kaida A, Harada H, Miura M. Radiosensitivity of quiescent and proliferating cells grown as multicellular tumor spheroids. Cancer science. 2017;108(4):704–12.

21. Raitanen J, Barta B, Hacker M, Georg D, Balber T, Mitterhauser M. Comparison of Radiation response between 2D and 3D cell culture models of different human Cancer cell lines. Cells. 2023;12(3):360.

22. Al-Ramadan A, Mortensen AC, Carlsson J, Nestor MV. Analysis of radiation effects in two irradiated tumor spheroid models. Oncology Letters. 2018;15(3):3008–16.

23. Menegakis A, Klompmaker R, Vennin C, Arbusà A, Damen M, van den Broek B, et al. Resistance of hypoxic cells to ionizing radiation is mediated in part via hypoxia-induced quiescence. Cells. 2021;10(3):610.

24. Cristini V, Koay E, Wang Z. An introduction to physical oncology: How mechanistic mathematical modeling can improve cancer therapy outcomes. CRC Press; 2017.

25. Deisboeck TS, Wang Z, Macklin P, Cristini V. Multiscale cancer modeling. Annual review of biomedical engineering. 2011;13(1):127–55.

26. Loessner D, Little JP, Pettet GJ, Hutmacher DW. A multiscale road map of cancer spheroids–incorporating experimental and mathematical modelling to understand cancer progression. Journal of cell science. 2013;126(13):2761–71.

27. Glazier J, Balter A, Poplawski N, Anderson A, Chaplain M, Rejniak K. Single-cell-based models in biology and medicine. Birkhauser-Verlag, Basel, Switzerland, chapter Magnetization to … ; 2007.

28. Wang Z, Butner JD, Kerketta R, Cristini V, Deisboeck TS. Simulating cancer growth with multiscale agent-based modeling. In: Seminars in cancer biology. vol. 30. Elsevier; 2015. p. 70–8.

29. Altrock PM, Liu LL, Michor F. The mathematics of cancer: integrating quantitative models. Nature Reviews Cancer. 2015;15(12):730–45.

30. Buch K, Peters T, Nawroth T, Sänger M, Schmidberger H, Langguth P. Determination of cell survival after irradiation via clonogenic assay versus multiple MTT Assay-A comparative study. Radiation oncology. 2012;7:1–6.

31. Rampersad SN. Multiple applications of Alamar Blue as an indicator of metabolic function and cellular health in cell viability bioassays. Sensors. 2012;12(9):12347–60.

32. Bonnier F, Keating ME, Wróbel TP, Majzner K, Baranska M, Garcia-Munoz A, et al. Cell viability assessment using the Alamar blue assay: A comparison of 2D and 3D cell culture models. Toxicology in Vitro. 2015 Feb;29(1):124–31. Available from: https://www.sciencedirect.com/science/article/pii/S0887233314001878.

33. Brüningk SC, Rivens I, Box C, Oelfke U, Ter Haar G. 3D tumour spheroids for the prediction of the effects of radiation and hyperthermia treatments. Scientific Reports. 2020;10(1):1653.

34. Kaida A, Miura M. Visualizing the effect of tumor microenvironments on radiation-induced cell kinetics in multicellular spheroids consisting of HeLa cells. Biochemical and Biophysical Research Communications. 2013;439(4):453–8.

35. Sharma M, Verma Y, Rao KD, Nair R, Gupta PK. Imaging growth dynamics of tumour spheroids using optical coherence tomography. Biotechnology Letters. 2007 Feb;29(2):273–8. Available from: https://link.springer.com/10.1007/s10529-006-9232-2.

36. Huang D, Swanson EA, Lin CP, Schuman JS, Stinson WG, Chang W, et al. Optical coherence tomography. science. 1991;254(5035):1178–81.

37. Azzollini S, Monfort T, Thouvenin O, Grieve K. Dynamic optical coherence tomography for cell analysis. Biomedical optics express. 2023;14(7):3362–79.

38. El-Sadek IA, Miyazawa A, Shen LTW, Makita S, Mukherjee P, Lichtenegger A, et al. Three-dimensional dynamics optical coherence tomography for tumor spheroid evaluation. Biomedical Optics Express. 2021 Nov;12(11):6844–63. Available from: https://opg.optica.org/boe/abstract.cfm?uri=boe-12-11-6844.

39. El-Sadek IA, Miyazawa A, Shen LTW, Makita S, Fukuda S, Yamashita T, et al. Optical coherence tomography-based tissue dynamics imaging for longitudinal and drug response evaluation of tumor spheroids. Biomedical Optics Express. 2020 Nov;11(11):6231–48. Available from: https://opg.optica.org/boe/abstract.cfm?uri=boe-11-11-6231.

40. Tan KH, Ang JLY, Yong ASK, Lim SZE, Kng JSJ, Liang K. Non-destructive viability assessment of cancer cell spheroids using dynamic optical coherence tomography with trypan blue validation. Biomedical Optics Express. 2024 Nov;15(11):6370–83. Available from: https://opg.optica.org/boe/abstract.cfm?uri=boe-15-11-6370.

41. Yang L, Yu X, Fuller AM, Troester MA, Oldenburg AL. Characterizing optical coherence tomography speckle fluctuation spectra of mammary organoids during suppression of intracellular motility. Quantitative Imaging in Medicine and Surgery. 2020;10(1):76.

42. Chhetri RK, Phillips ZF, Troester MA, Oldenburg AL. Longitudinal Study of Mammary Epithelial and Fibroblast Co-Cultures Using Optical Coherence Tomography Reveals Morphological Hallmarks of Pre-Malignancy. PLOS ONE. 2012 Nov;7(11):e49148. Available from: https://journals.plos.org/plosone/article?id=10.1371/journal.pone.0049148.

43. Monfort T, Azzollini S, Brogard J, Clémençon M, Slembrouck-Brec A, Forster V, et al. Dynamic full-field optical coherence tomography module adapted to commercial microscopes allows longitudinal in vitro cell culture study. Communications Biology. 2023;6(1):992.

44. Morishita R, Suzuki T, Mukherjee P, Abd El-Sadek I, Lim Y, Lichtenegger A, et al. Label-free intratissue activity imaging of alveolar organoids with dynamic optical coherence tomography. Biomedical Optics Express. 2023;14(5):2333–51.

45. Carmichael J, DeGraff WG, Gazdar AF, Minna JD, Mitchell JB. Evaluation of a tetrazolium-based semiautomated colorimetric assay: assessment of radiosensitivity. Cancer research. 1987;47(4):943–6.

46. Kim WH, Chon CY, Moon YM, Kang JK, Park IS, Choi HJ. Effect of anticancer drugs and desferrioxamine in combination with radiation on hepatoma cell lines. Yonsei medical journal. 1993;34(1):45–56.

47. Chen K, Swanson S, Bizheva K. Line-field dynamic optical coherence tomography platform for volumetric assessment of biological tissues. Biomedical Optics Express. 2024;15(7):4162–75.

48. Bratosin D, Mitrofan L, Palii C, Estaquier J, Montreuil J. Novel fluorescence assay using calcein-AM for the determination of human erythrocyte viability and aging. Cytometry Part A: the journal of the International Society for Analytical Cytology. 2005;66(1):78–84.

49. Crowley LC, Scott AP, Marfell BJ, Boughaba JA, Chojnowski G, Waterhouse NJ. Measuring Cell Death by Propidium Iodide Uptake and Flow Cytometry. Cold Spring Harbor protocols. 2016;2016(7).

50. Swanson S, Chen K, Cheraghi E, Osei E, Bizheva K. Longitudinal investigation of prostate tumor spheroid proliferation with dynamic line-field optical coherence tomography. bioRxiv; 2025. ISSN: 2692-8205 Pages: 2025.11.04.686594 Section: New Results. Available from: https://www.biorxiv.org/content/10.1101/2025.11.04.686594v1.

51. Thouvenin O, Boccara C, Fink M, Sahel J, Pâques M, Grieve K. Cell motility as contrast agent in retinal explant imaging with full-field optical coherence tomography. Investigative ophthalmology & visual science. 2017;58(11):4605–15.

52. Münter M, Vom Endt M, Pieper M, Casper M, Ahrens M, Kohlfaerber T, et al. Dynamic contrast in scanning microscopic OCT. Optics letters. 2020;45(17):4766–9.

53. Swat MH, Thomas GL, Belmonte JM, Shirinifard A, Hmeljak D, Glazier JA. Multi-scale modeling of tissues using CompuCell3D. In: Methods in cell biology. vol. 110. Elsevier; 2012. p. 325–66.

54. Gupta N, Lamborn K, Deen DF. A statistical approach for analyzing clonogenic survival data. Radiation research. 1996;145(5):636–40.

55. Srinivas US, Tan BW, Vellayappan BA, Jeyasekharan AD. ROS and the DNA damage response in cancer. Redox biology. 2019;25:101084.

56. Kim W, Lee S, Seo D, Kim D, Kim K, Kim E, et al. Cellular stress responses in radiotherapy. Cells. 2019;8(9):1105.

57. Wang JS, Wang HJ, Qian HL. Biological effects of radiation on cancer cells. Mil Med Res. 2018 Jun;5(1):20.

58. Lomax ME, Folkes LK, O’Neill P. Biological Consequences of Radiation-induced DNA Damage: Relevance to Radiotherapy. Clinical Oncology. 2013 Oct;25(10):578–85. Available from: https://www.clinicaloncologyonline.net/article/S0936-6555(13)00247-1/fulltext.

59. Reynolds P, Anderson JA, Harper JV, Hill MA, Botchway SW, Parker AW, et al. The dynamics of Ku70/80 and DNA-PKcs at DSBs induced by ionizing radiation is dependent on the complexity of damage. Nucleic acids research. 2012;40(21):10821–31.

60. Schmid TE, Dollinger G, Beisker W, Hable V, Greubel C, Auer S, et al. Differences in the kinetics of γ-H2AX fluorescence decay after exposure to low and high LET radiation. International journal of radiation biology. 2010;86(8):682–91.

61. O’Connell MJ, Walworth NC, Carr AM. The G2-phase DNA-damage checkpoint. Trends in cell biology. 2000;10(7):296–303.

62. Rainey M, Black E, Zachos G, Gillespie D. Chk2 is required for optimal mitotic delay in response to irradiation-induced DNA damage incurred in G2 phase. Oncogene. 2008;27(7):896–906.

63. Lauber K, Ernst A, Orth M, Herrmann M, Belka C. Dying cell clearance and its impact on the outcome of tumor radiotherapy. Frontiers in oncology. 2012;2:116.

64. Rajaee Z, Khoei S, Mahdavi SR, Ebrahimi M, Shirvalilou S, Mahdavian A. Evaluation of the effect of hyperthermia and electron radiation on prostate cancer stem cells. Radiation and environmental biophysics. 2018;57:133–42.

65. Cho YM, Kim YS, Kang MJ, Farrar WL, Hurt EM. Long-term recovery of irradiated prostate cancer increases cancer stem cells. The Prostate. 2012;72(16):1746–56.

66. Fazeli GR, Khouei S, Nikoufar A, Goliaei B. Reduced DNA damage in tumor spheroids compared to monolayer cultures exposed to ionizing radiation. INTERNATIONAL JOURNAL OF RADIATION RESEARCH. 2007.

67. Koch J, Moench D, Maass A, Gromoll C, Hehr T, Leibold T, et al. Three dimensional cultivation increases chemo-and radioresistance of colorectal cancer cell lines. PLoS One. 2021;16(1):e0244513.

68. Durand RE, Sutherland RM. Dependence of the radiation response of an in vitro tumor model on cell cycle effects. Cancer Research. 1973;33(2):213–9.

69. Horsman MR, Overgaard J. The impact of hypoxia and its modification of the outcome of radiotherapy. Journal of radiation research. 2016;57(S1):i90–8.

70. Jordan BF, Sonveaux P. Targeting tumor perfusion and oxygenation to improve the outcome of anticancer therapy. Frontiers in pharmacology. 2012;3:94.

71. Rey S, Schito L, Koritzinsky M, Wouters BG. Molecular targeting of hypoxia in radiotherapy. Advanced Drug Delivery Reviews. 2017;109:45–62.

72. Liu H, Adler AS, Segal E, Chang HY. A transcriptional program mediating entry into cellular quiescence. PLoS Genetics. 2007;3(6):e91.

73. Vinci M, Gowan S, Boxall F, Patterson L, Zimmermann M, Court W, et al. Advances in establishment and analysis of three-dimensional tumor spheroid-based functional assays for target validation and drug evaluation. BMC Biology. 2012 Mar;10(1):29. Available from: 10.1186/1741-7007-10-29.

74. Mittler F, Obeïd P, Rulina AV, Haguet V, Gidrol X, Balakirev MY. High-content monitoring of drug effects in a 3D spheroid model. Frontiers in oncology. 2017;7:293.

75. Elmore S. Apoptosis: a review of programmed cell death. Toxicologic pathology. 2007;35(4):495–516.

76. Elliott MR, Ravichandran KS. The dynamics of apoptotic cell clearance. Developmental cell. 2016;38(2):147–60.

77. Lammering G, Valerie K, Lin PS, Hewit TH, Schmidt-Ullrich RK. Radiation-induced activation of a common variant of EGFR confers enhanced radioresistance. Radiotherapy and oncology. 2004;72(3):267–73.

78. Xu W, Luo T, Li P, Zhou C, Cui D, Pang B, et al. RGD-conjugated gold nanorods induce radiosensitization in melanoma cancer cells by downregulating αvβ3 expression. International Journal of Nanomedicine. 2012:915–24.

79. Schweigerer L, Rave-Fränk M, Schmidberger H, Hecht M. Sublethal irradiation promotes invasiveness of neuroblastoma cells. Biochemical and biophysical research communications. 2005;330(3):982–8.

80. Hamis S, Kohandel M, Dubois LJ, Yaromina A, Lambin P, Powathil GG. Combining hypoxia-activated prodrugs and radiotherapy in silico: Impact of treatment scheduling and the intra-tumoural oxygen landscape. PLoS computational biology. 2020;16(8):e1008041.

81. Mao X, McManaway S, Jaiswal JK, Patel PB, Wilson WR, Hicks KO, et al. An agent-based model for drug-radiation interactions in the tumour microenvironment: Hypoxia-activated prodrug SN30000 in multicellular tumour spheroids. PLoS computational biology. 2018;14(10):e1006469.

82. Sánchez-Reyes A. A simple model of radiation action in cells based on a repair saturation mechanism. Radiation research. 1992;130(2):139–47.

83. Stewart RD. Two-lesion kinetic model of double-strand break rejoining and cell killing. Radiation research. 2001;156(4):365–78.

84. Cleri F. Agent-based model of multicellular tumor spheroid evolution including cell metabolism. The European Physical Journal E. 2019;42:1–15.

85. Kempf H, Hatzikirou H, Bleicher M, Meyer-Hermann M. In silico analysis of cell cycle synchronisation effects in radiotherapy of tumour spheroids. PLoS computational biology. 2013;9(11):e1003295.

86. Kempf H, Bleicher M, Meyer-Hermann M. Spatio-temporal dynamics of hypoxia during radiotherapy. PLoS One. 2015;10(8):e0133357.

87. Powathil GG, Adamson DJ, Chaplain MA. Towards predicting the response of a solid tumour to chemotherapy and radiotherapy treatments: clinical insights from a computational model. PLoS computational biology. 2013;9(7):e1003120.

88. Powathil GG, Swat M, Chaplain MA. Systems oncology: towards patient-specific treatment regimes informed by multiscale mathematical modelling. In: Seminars in cancer biology. vol. 30. Elsevier; 2015. p. 13–20.

89. Powathil GG, Munro AJ, Chaplain MA, Swat M. Bystander effects and their implications for clinical radiation therapy: Insights from multiscale in silico experiments. Journal of theoretical biology. 2016;401:1–14.

90. Thavarajah R, Mudimbaimannar VK, Elizabeth J, Rao UK, Ranganathan K. Chemical and physical basics of routine formaldehyde fixation. Journal of Oral and Maxillofacial Pathology : JOMFP. 2012;16(3):400–5. Available from: https://www.ncbi.nlm.nih.gov/pmc/articles/PMC3519217/.

