## Supplemental Model Information for "Comparison of volumetric dynamic optical coherence tomography with biological methods for evaluation of radiation effects in prostate tumor spheroids"

#### 1 Introduction

The particular form of agent-based modeling (ABM) used to simulate 3D tumor spheroid growth and radiation treatment is the cellular Potts model (CPM) [1]. Before detailing the implementation of the CPM, the mathematical theory of CPM is first given to provide context and notation. This document and its figures were adapted from Swanson [2].

#### 2 CPM theory

The mathematical formalism of the CPM described below arises directly from the Ising model description of a spin $\sigma$ located at the $i$-th position in a 2D or 3D lattice that is comprised of $N$ positions or voxels [3, 4].

##### 2.1 Lattice

Rather than spins, the lattice of the CPM is occupied by cells that each behave as an individual agent. Each unique cell is composed of multiple voxels that are assigned the same cell index $\sigma$. The surrounding environment or *Medium* is described by $\sigma = 0$, so in a simulation that contains $n$ cells, $\sigma$ belongs to the set $\{0, 1, ..., n\}$. The volume $v$ of the cell with index $\sigma^*$ is given in voxels by:

$$v(\sigma^*) = \sum_{i=0}^{N} \delta(\sigma_i, \sigma^*) \tag{1}$$

where $\sigma_i$ denotes the cell index at the $i$-th position in the lattice and $\delta$ denotes the Dirac delta function. Similarly, the surface area $s$ of cell $\sigma^*$ is calculated in pixels using the interfaces along adjacent cells as follows:

$$s(\sigma^*) = \frac{1}{2} \sum_{i,j \text{ neighbors}} \delta(\sigma_i, \sigma^*) \times [1 - \delta(\sigma_i, \sigma_j)] \tag{2}$$

While the cells are internally structureless, the resolution provided by spanning multiple lattice sites enables simulated cell behavior. Cells are grouped into biological cell types $\tau$ that have unique type-based attributes and behaviors, allowing for specific categorical phenotypes and morphologies. In addition to discrete cells that occupy the lattice, chemical fields governed by continuous partial differential equations (PDE)s may be discretized to the lattice to describe any fluid or chemical of interest.

#### 2.2 Modified Metropolis algorithm

The spatio-temporal behavior of each chemical field is dictated by its PDE. In contrast, the discrete cellular arrangement in the lattice evolves in time through a stochastic modified Metropolis algorithm [5], which is a Markov chain Monte Carlo method that performs "copy attempts" using the Hamiltonian to represent the effective energy of the lattice (Fig. 1). At the start of every copy attempt, a lattice site $i$ and a neighboring site $j$ are chosen at random with uniform distribution. The cell index $\sigma_j$ of the neighboring lattice site $j$ is copied into position $i$, then the effective energy before the copy attempt $\mathcal{H}$ is compared with the new effective energy $\mathcal{H}(\sigma_i = \sigma_j)$. A copy attempt is energetically favorable or unfavorable if the change in energy $\Delta\mathcal{H} = \mathcal{H}(\sigma_i = \sigma_j) - \mathcal{H}$ is negative or positive, respectively. Energetically favorable copy attempts are accepted, whereas energetically unfavorable copy attempts are accepted with a Boltzmann probability [6] as follows:

$$P(\sigma_i \to \sigma_j) = \begin{cases} 1 & \Delta\mathcal{H} \leq 0 \\ \exp(-\Delta\mathcal{H}/T) & \Delta\mathcal{H} > 0 \end{cases} \tag{3}$$

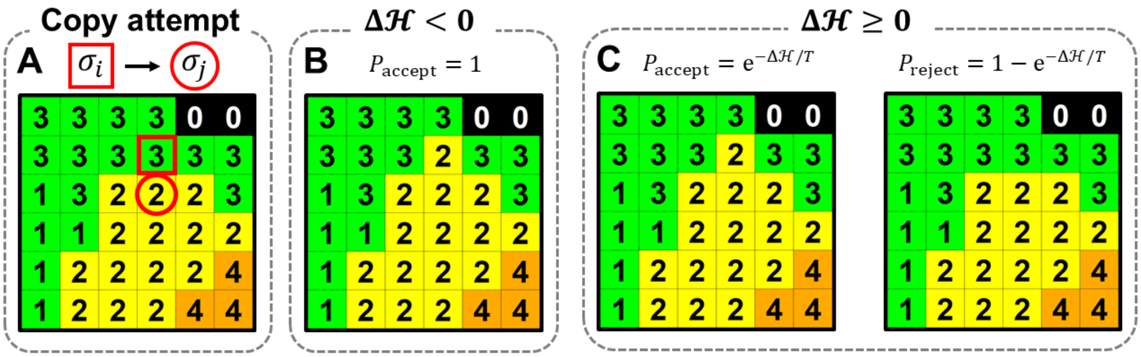


**Figure 1.** A CPM lattice with cell type $\tau$ and index $\sigma$ indicated by color and number, respectively. The depicted lattice has three cell types (green, yellow, and orange) in addition to the black *Medium*, including two unique cells $\sigma = 1$ and $\sigma = 3$ of different volumes but the same green type. A) The modified Metropolis algorithm performs "copy attempts" by randomly selecting a lattice site $i$ (in red square) and a neighboring site $j$ (in red circle), and proposing that the cell index $\sigma$ at $i$ becomes that of $j$. B) The change is accepted if the change in effective energy $\Delta\mathcal{H}$ is reduced. C) Otherwise, the change is accepted with probability $\exp(-\Delta\mathcal{H}/T)$.

Another remanent of its origin in statistical physics, $T$ is called the temperature of the model, although here it has units of simulated energy. $T$ is also called the amplitude of cell-membrane fluctuations or the cell motility because each successful copy attempt changes a single lattice site along the border of one cell – or two cells if the interface falls along another cell border rather than the medium. Periodic boundary conditions are often imposed to the lattice, and may be thought to map the simulation space onto a torus. That is, cells that fluctuate beyond the edge of the cell lattice appear to enter the lattice on the opposite edge. However, fixed boundary conditions of a confined environment can also be imposed. One Monte Carlo step (MCS) has passed when $N$ copy attempts have been performed on a lattice of $N$ voxels. As such, the MCS demarcates the passage of simulated time.

Note that the ratio $\Delta\mathcal{H}/T$ in Eq. 3 controls how frequently cells rearrange. Large $\Delta\mathcal{H}/T$ results in rigid, barely motile cells whereas small $\Delta\mathcal{H}/T$ results in high cell motility and rearrangement. Consequently, simulations that impose a higher temperature $T$ parameter observe higher rates of cellular rearrangement. The nature of those rearrangements, however, is directed by the form of the Hamiltonian $\mathcal{H}$, which embodies the physical and phenomenological forces enacted on the cells.

#### 2.3 Hamiltonian

The Hamiltonian is the effective energy of the simulation space through which all rules that dictate cellular characteristics and intercellular interactions are ultimately manifested via regulation of cell surface fluctuations. Common components of the CPM Hamiltonian that are relevant here include contributions from intercellular contact adhesion, cell size, and external forces like gravity:

$$\mathcal{H} = \mathcal{H}_a + \mathcal{H}_v + \mathcal{H}_s + \mathcal{H}_g \tag{4}$$

The following sections describe the mathematical form of each of these contributions.

##### 2.3.1 Contact adhesion

The contact adhesion term $\mathcal{H}_a$ describes the energy associated with cell-cell interfaces of adjacent cells through a matrix $J$ that is indexed by cell type $\tau$ as follows:

$$\mathcal{H}_a = \sum_{i,j \text{ neighbors}} J\{\tau(\sigma_i), \tau(\sigma_j)\} \times [1 - \delta(\sigma_i, \sigma_j)] \tag{5}$$

Thus, the effective energy contribution from contact adhesion sums the cell type specific contact energy between all pairs of neighboring lattice sites $i$ and $j$ that belong to different cells. Contact adhesion models surface tension because $J\{\tau(\sigma_i), \tau(\sigma_j)\}$ is generally positive, in which case the Metropolis algorithm (Eq. 3) minimizes the surface area of cells that is shared with other cells or the medium [7].

##### 2.3.2 Cell size

The size of cells is typically determined by assigning each cell a target volume $V_t$ and imposing the following:

$$\mathcal{H}_v = \sum_{\sigma=0}^{n} \lambda_v(\sigma_i) \times [v(\sigma_i) - V_t(\sigma_i)]^2 \tag{6}$$

where the cell volume $v$ adheres to its target more strictly with increasing $\lambda_v$. Similarly, the cell surface area $s$ is encouraged to achieve a target surface area $S_t$ with strictness $\lambda_s$ as follows:

$$\mathcal{H}_s = \sum_{\sigma=0}^{n} \lambda_s(\sigma_i) \times [s(\sigma_i) - S_t(\sigma_i)]^2 \tag{7}$$

##### 2.3.3 Gravity

Gravity is implemented as follows:

$$\mathcal{H}_g = \sum_{i=0}^{N} \lambda_g(z_{\max} - z_i) \times [1 - \delta(\sigma_i, 0)] \tag{8}$$

where $z_i$ is the distance along the $z$-axis of position $i$ with maximum distance $z_{\max}$ and $\lambda_g$ mediates the strength of the gravitational field. Notice that only cells experience the gravitational force, with the *Medium* excluded from the calculation.

#### 3 CPM implementation

Using the CompuCell3D (CC3D) platform [8], radiation treatment was incorporated into a CPM that models prostate tumor spheroid growth via the 3D Petri Dish method. It directly employs the Hamiltonian given by Eq. 4.

In addition to the *Medium* and *Gel* cell types that describe the culture medium and agarose gel, respectively, there are three tumor cell types: *Normoxic*, *Hypoxic*, and *Necrotic*. These tumor cell types are updated every 10 MCS using the partial pressure of oxygen $P_O$ at the centre of mass (COM) of the cell $i_{\text{COM}}$. A tumour cell is *Normoxic*, *Hypoxic*, or *Necrotic* if its partial pressure of oxygen is respectively above, between, or below the hypoxic and necrotic oxygen thresholds of 5 mmHg and 1 mmHg. While transitions between the *Normoxic* and *Hypoxic* cell types are reversible, conversion to the *Necrotic* type indicates irreversible cell death due to starvation.

Oxygen is maintained at a constant partial pressure in *Medium* and modeled with a reaction-diffusion PDE as follows:

$$\frac{\partial P_O(i,t)}{\partial t} = D_O \nabla^2 P_O(i,t) - f_{\text{uptake}}(i,t) \times \delta(\tau(\sigma_i), \tau_{\text{tum}}) \tag{9}$$

where the diffusion coefficient $D_O$ is 3440 pixel$^2$/MCS and roughly corresponds to 1200 $\mu$m$^2$/s, which is slightly more than half the diffusion constant of oxygen in water [9] under the assumption that the physical size of each simulated voxel is (5 $\mu$m)$^3$ or 125 $\mu$m$^3$. The function $f_{\text{uptake}}$ describes the consumption of oxygen per MCS within living *Normoxic* and *Hypoxic* tumor cell types, denoted collectively as $\tau_{\text{tum}}$, through the following piecewise linear approximation to a Michaelis-Menten saturation curve:

$$f_{\text{uptake}}(i,t) = \begin{cases} \alpha P_O(i,t) & P_O(i,t) < P_{\text{O, max}} \\ \alpha P_{\text{O, max}} & P_O(i,t) \geq P_{\text{O, max}} \end{cases} \tag{10}$$

Here, $\alpha$ is the fraction of the local oxygen that is consumed until saturation at the maximum partial pressure $P_{\text{O, max}}$.

In conjunction with consumption, the partial pressure of oxygen at the COM of each living cell $i_{\text{COM}}$ also increases its target volume $V_t$ via the following Michaelis-Menten saturation curve:

$$\frac{dV_t}{dt} = \frac{V_{\text{max}} \times P_O(i_{\text{COM}}, t)}{K_V + P_O(i_{\text{COM}}, t)} \tag{11}$$

where $V_{\text{max}}$ and $K_V$ denote the maximum growth rate and half saturation oxygenation, respectively. Similarly, the cell's target surface area $S_t$ grows with maximum growth rate $S_{\text{max}}$ and half saturation oxygenation $K_S$ as follows:

$$\frac{dS_t}{dt} = \frac{S_{\text{max}} \times P_O(i_{\text{COM}}, t)}{K_S + P_O(i_{\text{COM}}, t)} \tag{12}$$

Once *Normoxic* and *Hypoxic* cells have achieved a sufficient volume called the mitotic volume, they undergo mitosis and split along a random direction into two identical cells that are assigned post-mitotic target volumes and surface areas.

The simulation parameters were determined phenomenologically by comparison with quantitative measurement of the size and shape of the prostate tumor spheroid and its containing well inside the agarose gel via volumetric optical coherence tomography (OCT) imaging [10] (Fig. 2). These measurements also determined the temporal calibration of 1200 MCS to 24 hours, as we assume simulated and physical time are proportional [8]. To replicate the gaps observed throughout the spheroid in accordance with the CPM contact adhesion implementation, a phenomenological *Gap* cell type was developed to mimic pockets of culture medium trapped inside the spheroid. Radiation treatment was then implemented as described in the manuscript.

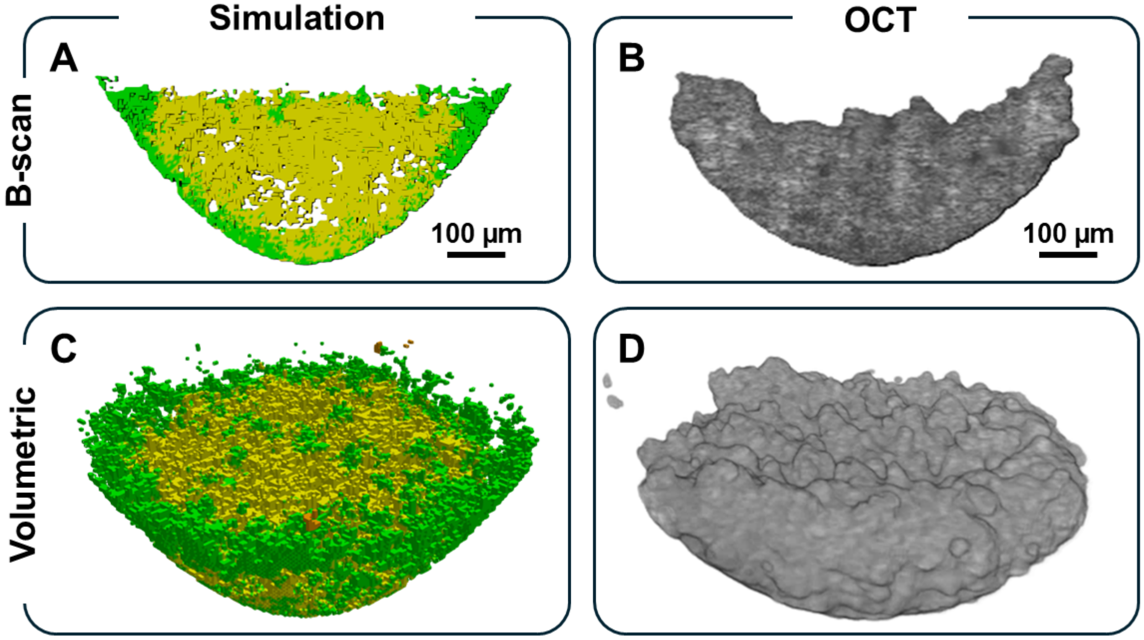


**Figure 2.** The simulation parameters were determined phenomenologically by comparing morphology with OCT imaging and masking of cultured prostate tumor spheroids. Cross-section B-scan (top row) and volumetric (bottom row) visualizations of a spheroid 48 hours after seeding generated by simulation (left column) and OCT (right column), with surrounding culture medium and gel removed. Scale bar: 100 $\mu$m.

### 4 Code availability

The full code of the CPMs modeling prostate tumor spheroid growth with and without radiation treatment are publicly available on GitHub at github.com/skswanso/CPM.git. A brief explanation of the files composing the CC3D platform follows. Each CPM is governed by a CC3D file, an XML file, a main Python file, and any number of Python files called "Steppables" that are implemented at a chosen frequency, given in number of MCS. The CC3D file simply points towards the locations of the other modeling files, relative to its own file location. The XML file of each model contains specific parameters required by CC3D, and has been extended here to further centralize and store all parameter values, which are accessed by the relevant Python file. The control of each Steppable python file is managed through the main Python file. The Growth Steppable controls conversion between cell types and implements cell growth by increasing cell target volumes and surface areas. The Mitosis Steppable manages cell division. The Output Steppable collects, calculates and manages storage of output data. Finally, the CPM of radiation treatment additionally includes a Radiation Steppable that identifies cells that survive treatment.

### References


1. Graner F, Glazier JA. Simulation of biological cell sorting using a two-dimensional extended Potts model. Physical review letters. 1992;69(13):2013.
2. Swanson S. Characterization of prostate tumor spheroid growth and response to treatment using dynamic optical coherence tomography [PhD thesis]. Waterloo, Canada: University of Waterloo; 2025. Available at `https://hdl.handle.net/10012/22193`.
3. Ising E. Beitrag zur theorie des ferromagnetismus. Zeitschrift für Physik. 1925;31(1):253-8.

4. Lenz W. Beitrag zum Verständnis der magnetischen Erscheinungen in festen Körpern. Z Phys. 1920;21:613-5.

5. Cipra BA. An introduction to the Ising model. The American Mathematical Monthly. 1987;94(10):937-59.

6. Metropolis N, Rosenbluth AW, Rosenbluth MN, Teller AH, Teller E. Equation of state calculations by fast computing machines. The journal of chemical physics. 1953;21(6):1087-92.

7. Glazier JA, Graner F. Simulation of the differential adhesion driven rearrangement of biological cells. Physical Review E. 1993;47(3):2128.

8. Swat MH, Thomas GL, Belmonte JM, Shirinifard A, Hmeljak D, Glazier JA. Multi-scale modeling of tissues using CompuCell3D. In: Methods in cell biology. vol. 110. Elsevier; 2012. p. 325-66.

9. Shirinifard A, Gens JS, Zaitlen BL, Popławski NJ, Swat M, Glazier JA. 3D multi-cell simulation of tumor growth and angiogenesis. PloS one. 2009;4(10):e7190.

10. Swanson S, Chen K, Cheraghi E, Osei E, Bizheva K. Longitudinal investigation of prostate tumor spheroid proliferation with dynamic line-field optical coherence tomography. bioRxiv; 2025. ISSN: 2692-8205 Pages: 2025.11.04.686594 Section: New Results. Available from: https://www.biorxiv.org/content/10.1101/2025.11.04.686594v1.
